# IDEA: a web server for Interactive Differential Expression Analysis with R Packages

**DOI:** 10.1101/360461

**Authors:** Qi Zhao, Yubin Xie, Peng Nie, Rucheng Diao, Lichen Sun, Zhixiang Zuo, Jian Ren

## Abstract

Differential expression (DE) analysis is a fundamental task in the downstream analysis of the next-generation sequencing (NGS) data. Up to now, a number of R packages have been developed for detecting differentially expressed genes. Although R language has an interaction-oriented programming design, for many biology researchers, a lack of basic programming skills has greatly hindered the application of these R packages. To address this issue, we developed the Interactive Differential Expression Analyzer (IDEA), a Shiny-based web application integrating the differential expression analysis related R packages into a graphical user interface (GUI), allowing users to run the analysis without writing any new code. A wide variety of charts and tables are generated to facilitate the interpretation of the results. In addition, IDEA also provides a combined analysis framework which helps to reconcile any discrepancy from different computational methods. As a public data analysis server, IDAE is implemented in HTML, CSS and JavaScript, and is freely available at http://idea.renlab.org.

## Introduction

High-throughput sequencing technology is rapidly becoming the standard method for measuring gene expression at the transcriptional level. One of the major goals of such work is to identify differentially expressed (DE) genes under two or more conditions. A number of computational tools, such as DESeq2 [1], edgeR [2], NOISeq [3], PoissonSeq [4], SAMseq [5] and Cuffdiff [6] have been developed for the analysis of differential gene expression from RNA-seq data. Most of these tools are implemented in R language, which is commonly used for the analysis of high-dimensional expression data. However, a fairly high level of programing skill is required when applying these R tools to screen out differentially expressed genes, which greatly hinders the application of these tools since many biology researchers have little programing experience. In recent year, a number of software, such as easyRNASeq[7], RNASeqGUI[8] and RobiNA[9] et al, was developed to address this issue. Although the above tools have provided convenient GUI-based computer platforms for back-end R packages to some extent, none of them implements an interactive interface that can not only facilitate the understanding of analysis process for a web-bench scientist, but also inconvenient to adjust the analytical parameters, even for advanced users. Moreover, since different packages will generate inconsistent results, it is difficult for the users to decide which DE algorithm to use. Thus, an interactive platform that can combine these tools together is necessary for obtaining more solid analysis results.

In this regards, we present the Interactive Differential Expression Analyzer (IDEA), a Shiny-based web application dedicated to the identification of differential expression genes in an interactive way. IDEA was built as a user-friendly and highly interactive utility using the Shiny package in R. Besides, five relevant R packages are integrated into IDEA. IDEA is capable of visualizing the results with plenty of charts and tables, as well as providing great ease of interaction during the analysis

## Features and options

### Framework

IDEA provides a well-designed data analysis framework to help identify reliable differential expressed features (Figure 1, 2A). Briefly, three kinds of experimental comparison between groups can be performed using IDEA, namely “standard comparison”, “multifactor design” and “comparison without replicates”. To have a quick start, IDEA requires users to specify the experimental design and can automatically determine available packages for further analysis. For input dataset, a read count matrix and a design matrix which presents the relationship between samples and conditions were mandatory for an independent analysis. Users were encouraged to load feature length file to obtain length-normalized expression values, such as (Reads Per Kilobase Million) RPKM and (Fragments Per Kilobase Million) FPKM. After that, data exploration panel was introduced to help assessing the data quality and normalizing the reads count dataset for further analysis. Differential expression analysis can be sequentially selected for running separately or not. As different algorithms may generate inconsistent differential expression feature list, we further developed a combine analysis module in IDEA to integrate result from each selected analysis packages. Additionally, a build-in example data that enabled users to run demo analysis and download demo report was established.

**Figure 1.**
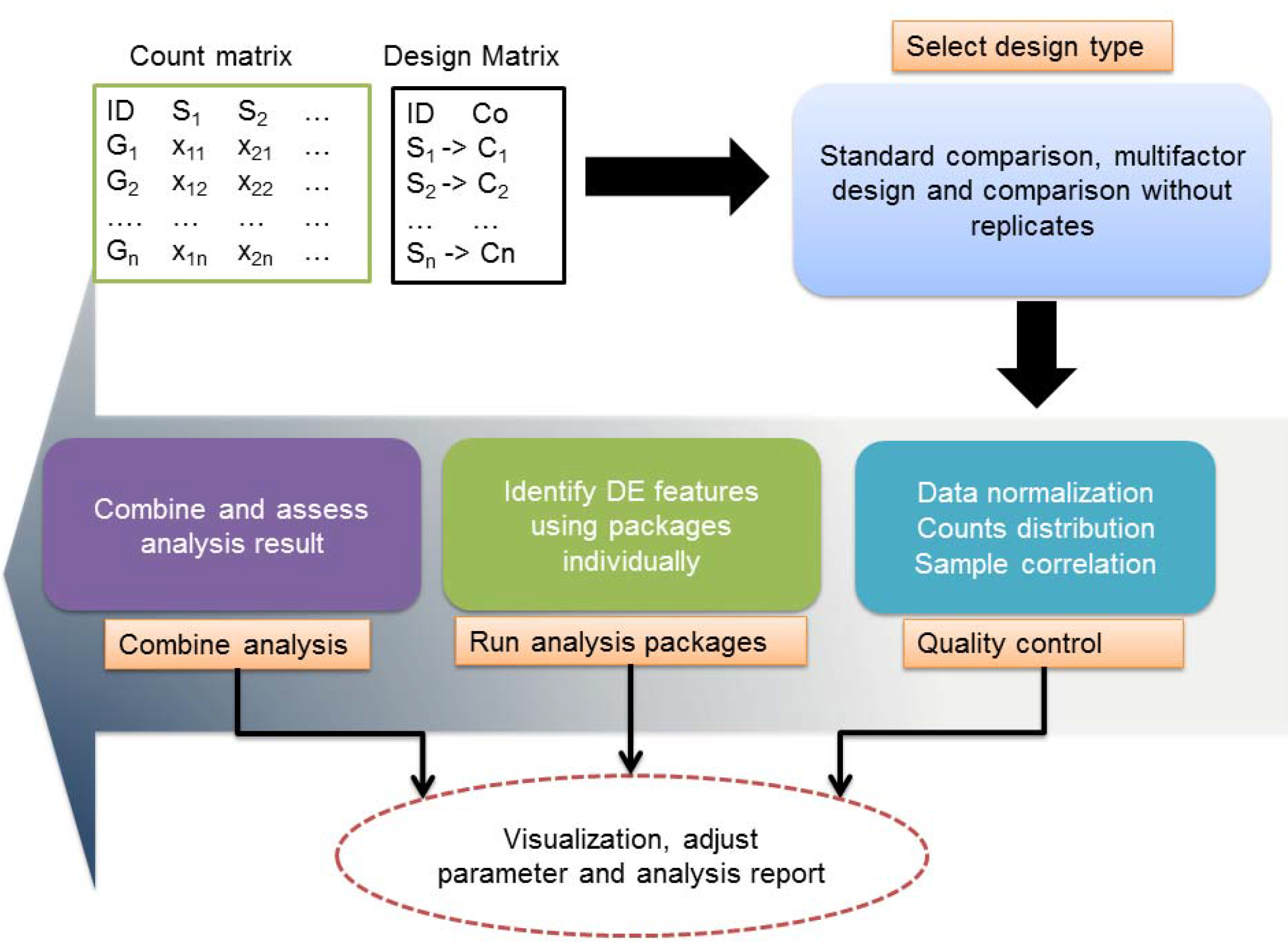
Differential expression analysis framework implemented in IDEA.

**Figure 2.**
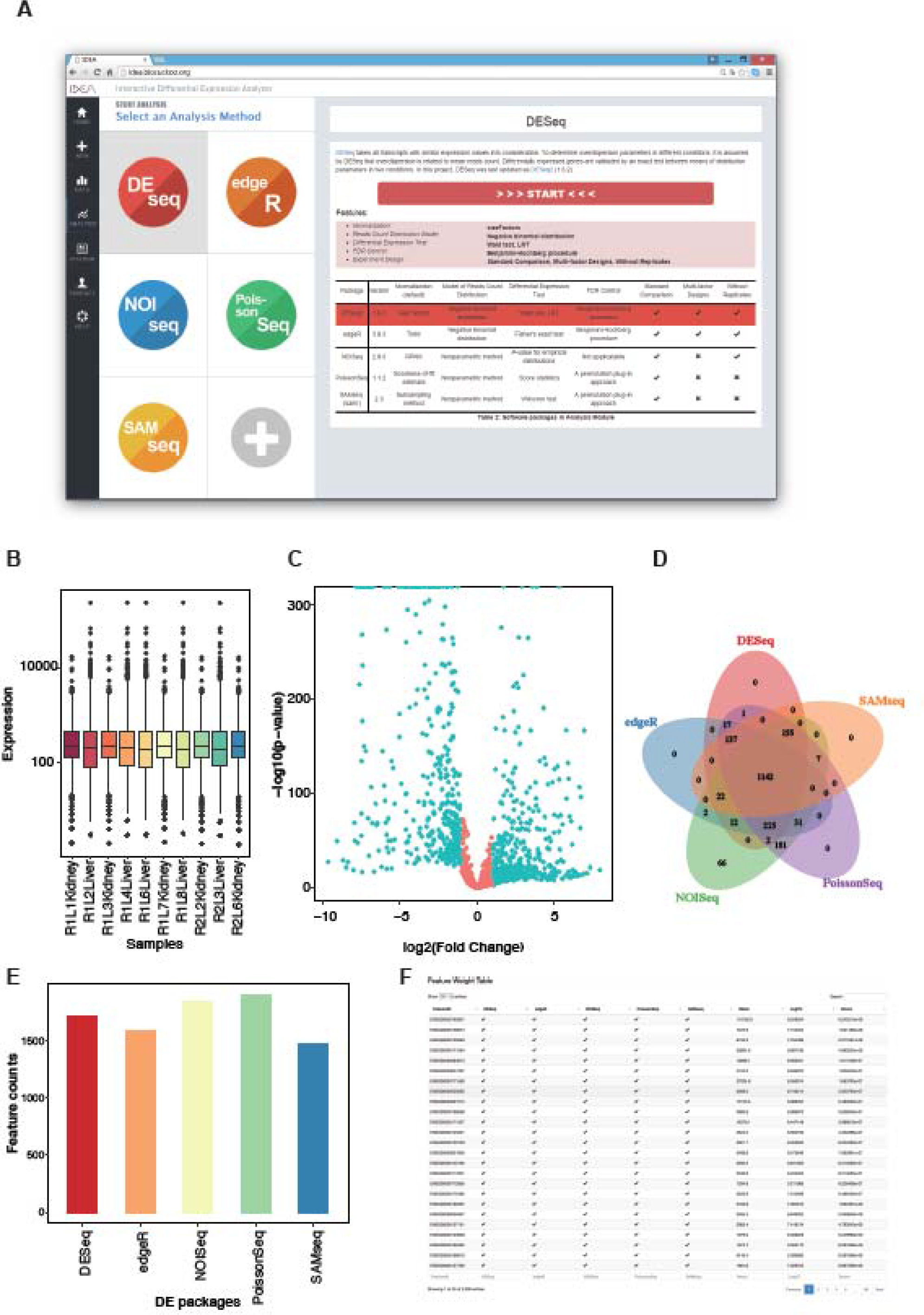
Selected analysis plots generated by IDEA. A, screen shot of IDEA web page. B, a box plot presents the normalized reads count distribution among samples. C, Differential expression analysis result was shown in a volcano plot; each point represents a feature for statistic test, the green points are significantly differential expressed with adjusted P value < 0.05. D, Overlap analysis based on the DE result from five analysis packages. E, overview of differential expressed feature count by each package shown in a bar plot. F. combine analysis result data frame, only top 50 features were displayed as default.

#### Data exploration

Data quality assessment and quality control are essential steps for expression analysis. Beyond the reads base quality and mapping quality evaluation in raw data preprocessing stage, IDEA implements a quality control panel based on reads count distribution in terms of genes and samples. In this panel, we provide several normalized methods including TPM, upper quartile (Default), RPKM and “none” to help exploring potential calculating bias from different normalization approaches. We also provide three visualization strategies: *density area plot*, *bar plot* and *box plot* to help assessing the data quality (Figure 2B). These plots illustrate the feature expression distribution in each sample, allowing users to identify samples with extremely low or high expressed gene number.

Expression-based principal component analysis (PCA) and correlation analysis were provided for studying the sample relationship. *Scatter plot* with sample point and *heat map* presenting spearman correlation coefficients were generated respectively. Specifically, if the samples come from the same condition, we expect to observe a closer sample distance in the scatter plot, and to find the tested samples under the resembling branch within clustered heat map. Generally, biological or technical replicates exhibit higher correlation than samples from different sources, thus an extra correlation scatter plot containing detail relationships between couple of samples was included. It gives a further investigation of correlation analysis result from clustered heat map.

It’s also crucial for biologist to verify the known or some housekeeping genes in the experiment. By setting negative and positive control in a “wet” experiment, features with known expression patterns can be applied for assessing the quality of sequencing library, such as low expression of a gene in knockout library against wild type. IDEA, therefore, provides a *bar plot* interface to help users query the average abundance of a specified gene in each condition, while some internal control features can be checked with the help of this function.

#### Multiple DE methods integration

To allow users with little programming experience to easily perform DE analysis using the most commonly used R packages, the current implementation of IDEA has integrated DESeq2, edgeR, NOISeq, PossionSeq and SAMseq into a convenient web-based GUI. To avoid tedious operations, parameters of each method were optimized to fit most cases, and only few ones were required be adjusted in special scenario. When the analysis is finished, important information involved in each method was also presented, including model parameters estimation, normalized factor and statistic descriptions. Plots, like *MA plot*, *volcano plot* (Figure 2C), clustered *heat map* and *FDR*/*P-value distribution bar plot*, are presented to summarize the identified DE genes.

Different packages may generate discrepant results on the same dataset. To reconcile any such discrepancy, we developed a combination analysis module to integrate the results from the different packages in an unbiased manner. Using a set of rank aggregation methods [10], such as Robust Rank Aggregation and Stuart P-value Integration, the integration score of a given DE feature is calculated as the mean of the integrated rank numbers generated by each of the packages. To visualize the combined results, a *bar plot* showing the DE features identified by each packages (Figure 2D) and a Venn diagram demonstrating the distribution and reliability of the verified features (Figure 2E) were developed. Finally, the combined result was also presented in an interactive table to help exploring the most confidential DE features (Figure 2F).

### Interactive analysis and fancy analysis report

Interactive features involved in IDEA consist of dynamic data operation, parameter adjustment and real-time plot rendering. First, dynamic data table enables interactively filtering, sorting and querying genes from input and out data frames by sorts of attributes (Figure 3A). Second, parameter adjustment enables users to refine analysis with modified parameters in real time without rerun the irrelevant step. For example, in DEseq analysis section, modifying DE gene filtering criterion only require to trigger transformation of the result table, and the DEseq will not be executed again. Similarly, in combine analysis, modifying the parameters of one specific tool do not affect others. This may greatly reduce the reanalysis time for exploring analysis, and especially suitable for screening gene lists that are stable under different parameters. For interactive plot rendering, we added some switchable panels in plot frame for refining plot setting in real time. Samples, colors, comparisons and plot data can be dynamically adjusted as well (Figure 3B, 3C). Those options can just trigger re-execution of the plot function while leave the computing function unchanged. Plots can be interactive repainted while the corresponding parameters altered.

**Figure 3.**
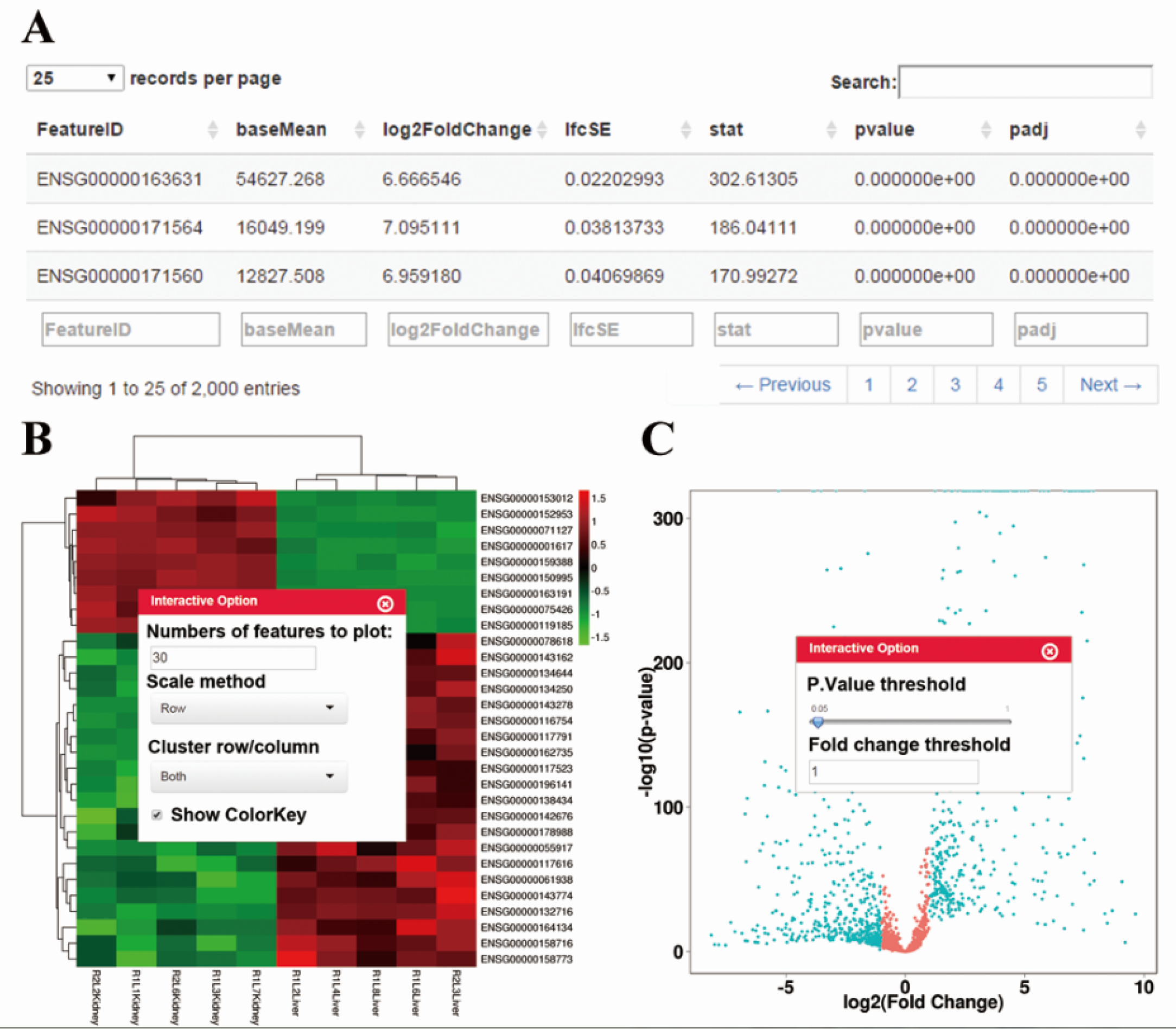
Interactive features provided in IDEA. A. dynamic data table JavaScript library was applied to help interactively exploring features stored in data frame. B. The floating control panel that enables modifying parameters in real-time.

Using R markdown and knitr package, IDEA summarized analysis results into a well-structured fancy report with all parameters, figures and citations listed. To meet the publication requirement, high quality figures with 300 dpi and result table can be downloaded from the final report, separately.

## Implementation

IDEA was initially implemented in R programming language [https://www.r-project.org/] with shiny [http://shiny.rstudio.com/] for building interactive web applications. It hosted in a Unix-like system which have both R and shiny server pre-installed. The front-end code was elaborately reconstructed with JavaScript, CSS and html to optimize the interactive experience of users. We also implemented an R package version of IDEA that allows users to run analysis locally under R environment. The package can be freely accessed and installed from https://github.com/likelet/IDEA. Detailed usage information was presented at the ‘help’ section of the web page.

## Future developments

As IDEA was developed under shiny web framework, its web utilities were greatly limited by shiny server performance. Therefore, to lift the limitation of server resources, we encourage users to install the local version when performing differential expression analysis. Besides, for interactive analysis, plots were dynamically rendered when data or parameters have changed, while a true sense of interactive plot have not yet been implemented due to a possible heavy browser loading. We will overcome this with reduced data points and customized data structure in the near future. Popular data visualization JavaScript library like D3 (https://d3js.org), plotly (https://plot.ly) and Highchart (https://www.highcharts.com) will be integrated as well. For the extension of more functions, extra analysis features, such as gene annotation, function enrichment and co-expression analysis, will be included into IDEA in the near future.

Nowadays, various third-party tools were developed to perform differential expression analysis [11], while some of them were not implemented in R. In the current version, IDEA only selected five most popular and efficient R-based tools in the analysis pipeline. For future development, packages in low computing efficiency will be integrated in R package version as an optional analysis module. However, non R-based tools like gfold[12], cufflinks[13] and Stringtie[14] etc, will not directly supported in our platform. Instead, to cope alternately with this issue, we are going to provide a combine analysis framework based on results generated by those tools.

## Conclusion

IDEA is a user-friendly web application for differential expression analysis, which combines the five commonly used DE analysis packages into a highly interactive web-based framework using the Shiny package. The abundant visualization functions in IDEA enable the users to gain both deeper insight into their original data and the ability to interpret the results in a more intuitive way. Furthermore, in contrast to the static image generated by R language, IDEA can produce diagrams in real time, which makes the analytical process more convenient and efficient. Taken together, we introduce IDEA as an effective application that can facilitate and simplify the process of differential expression analysis in high-dimension expression data.

